# Massively multiplex single-cell Hi-C

**DOI:** 10.1101/065052

**Authors:** Vijay Ramani, Xinxian Deng, Kevin L Gunderson, Frank J Steemers, Christine M Disteche, William S Noble, Zhijun Duan, Jay Shendure

## Abstract

We present combinatorial single cell Hi-C, a novel method that leverages combinatorial cellular indexing to measure chromosome conformation in large numbers of single cells. In this proof-of-concept, we generate and sequence combinatorial single cell Hi-C libraries for two mouse and four human cell types, comprising a total of 9,316 single cells across 5 experiments. We demonstrate the utility of single-cell Hi-C data in separating different cell types, identify previously uncharacterized cell-to-cell heterogeneity in the conformational properties of mammalian chromosomes, and demonstrate that combinatorial indexing is a generalizable molecular strategy for single-cell genomics.

## Main Text

Our understanding of genome architecture has largely progressed through the successive development of new technologies^1^. Advances in microscopy revealed the presence of “chromosome territories”—nuclear regions that preferentially self-associate in a manner correlated with transcriptional activity^2^. The invention of Chromosome Conformation Capture (3C) and its derivatives^3^ resulted in a proliferation of data measuring genome architecture and its relation to other aspects of nuclear biology at increasing resolution.

3C assays rely on the concept of proximity ligation, a technique that has been used to measure local protein-protein^4^, RNA-RNA^5^, and DNA-DNA interactions^6^. By coupling an “all-vs-all” 3C assay with massively parallel sequencing^7,8^ (*e.g.* “Hi-C”), one is able to query relative contact probabilities genome-wide. However, contact probabilities generated by these assays represent ensemble averages of the respective conformations of the millions of nuclei used as input, and scalable techniques characterizing the variance underlying these population averages remain largely underdeveloped. A pioneering study in 2013 demonstrated proof-of-concept that Hi-C could be performed on single isolated mouse nuclei, but relied on the physical separation and processing of single murine cells in independent reaction volumes, with consequent low-throughput^9^.

The repertoire of high-throughput single-cell techniques for other biochemical assays has expanded rapidly as of late^10-13^. Single-cell RNA-seq (scRNA-seq) was recently paired with droplet-based microfluidics to markedly increase its throughput^11,12^. Orthogonally, we introduced the concept of combinatorial cellular indexing^10^, a method that eschews microfluidic manipulation and instead tags the DNA within intact nuclei with successive (combinatorial) rounds of nucleic acid barcodes, to measure chromatin accessibility (scATAC-seq) in thousands of single cells without physically isolating each single cell. However, such throughput-boosting strategies have yet to be successfully adapted to single-cell chromosome conformation analysis.

To address this gap, we sought to develop a high-throughput, easy-to-implement single-cell Hi-C protocol (Figure 1a), based on the concept of combinatorial indexing and also building on recent improvements to the Hi-C protocol^14,15^. A population of cells is fixed, lysed to generate nuclei, and restriction digested *in situ* with the enzyme DpnII. Nuclei are then distributed to 96 wells, wherein the first barcode is introduced through ligation of barcoded biotinylated double-stranded bridge-adaptors. Intact nuclei are then pooled and proximity ligated all together, followed by dilution and redistribution to a second 96- well plate. Importantly, this dilution is carried out such that each well in this second plate contains at most 25 nuclei. Following lysis, a second barcode is introduced through ligation of barcoded Y-adapters.

**Figure 1:**
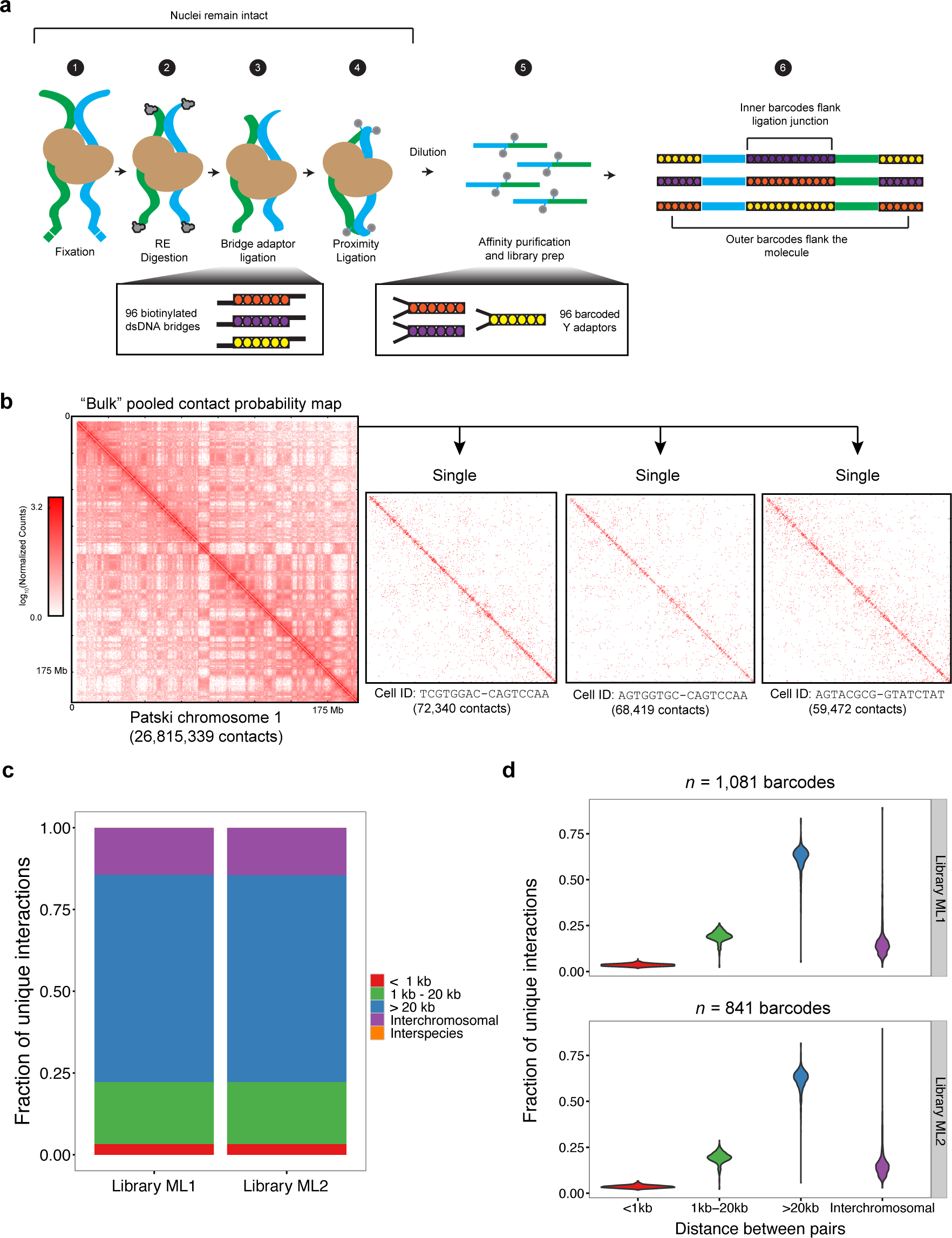
Combinatorial single cell Hi-C integrates the *in situ* Hi-C protocol with combinatorial cellular indexing to generate signal-rich bulk Hi-C maps that can be decomposed into single cell Hi-C maps. a.) Combinatorial single cell Hi-C follows the traditional paradigm of fixation, digestion, and re-ligation shared by all Hi-C assays (Steps 1 – 4), but importantly uses a biotinylated bridge adaptor to incorporate a first round of barcodes prior to proximity ligation (Step 3), and custom barcoded Illumina Y-adaptors (Step 5) to incorporate a second round of barcodes prior to affinity purification and library amplification (Steps 5 – 6). b.) Bulk data generated by this protocol can be decomposed to single cell Hi-C maps. Shown are combinatorial single cell Hi-C reads for mouse chromosome 1, corresponding to data from the Patski cell line. This data can then be separated on the basis of cellular indices into many single cell contact maps, shown here for three single Patski cells. c.) Our Hi-C libraries demonstrate a high *cis*:*trans* ratio, measured as the ratio of intrachromosomal contacts > 20 kb apart to interchromosomal contacts. d.) The high *cis*:*trans* ratio observed in bulk data is maintained after libraries are decomposed to ∼1800 single cell Hi-C maps. All indices tagging fewer than 1,000 reads are ignored in this analysis.

As the number of barcode combinations (96 x 96) exceeds the number of nuclei (96 x 25), the vast majority of single nuclei are tagged by a unique combination of barcodes. All material is once again pooled, and biotinylated junctions are purified with streptavidin beads, restriction digested, and further processed to Illumina sequencing libraries. Sequencing these molecules with relatively long paired-end reads (*i.e.* 2 x 250 base pair (bp)) allows one to identify not only the genome-derived fragments of conventional Hi-C, but also external and internal barcodes (each combination of which is hereafter referred to as a ‘cellular index’) which enable decomposition of the Hi-C data into single-cell contact probability maps (Figure 1b). Like scATAC-seq with combinatorial cellular indexing^10^, this protocol can process hundreds to thousands of cells per experiment without requiring the physical isolation of each cell.

As a proof-of-concept, we applied combinatorial single cell Hi-C to synthetic mixtures of cell lines derived from mouse (primary mouse embryonic fibroblasts (MEFs), and the ‘Patski’ embryonic fibroblast line) and human (HeLa S3, the HAP1 cell line, K562, and GM12878; all five experiments and sequenced libraries are summarized in Table 1, although we focus on ML1 and ML2 biological replicates in the text). All experiments were carried out such that subsets of cell types received specific barcodes during the first round of barcoding ( *e.g.* in ML1 and ML2, each well during the first round of barcoding contained either HeLa S3 + Patski cells or HAP1 + MEF cells; see **Methods**).

**Table 1:** Summary of all 5 sequenced libraries in this study.

|  | **ML1** | **ML2** | **PL1** | **PL2** | **ML3** |
| --- | --- | --- | --- | --- | --- |
| **Programming During First Barcoding / Cell Type Composition** | HeLa S3 + Patski mixed in 4 rows; HAP1 + MEF mixed in 4 rows | HeLa S3 + Patski mixed in 4 rows; HAP1 + MEF mixed in 4 rows | HeLa S3 in 2 rows; Patski in 2 rows; HAP1 in 2 rows; MEF in 2 rows | HeLa S3 in 2 rows; Patski in 2 rows; HAP1 in 2 rows; MEF in 2 rows | GM12878 + Patski mixed in 4 rows; K562 + MEF mixed in 4 rows |
| **# of sequenced reads** | 128,660,854 | 124,833,616 | 107,609,682 | 181,089,598 | 20,302,425 |
| **# of associated reads** | 62,566,086 | 59,931,225 | 59,054,337 | 94,661,167 | 10,192,548 |
| **# of mapped, associated, MAPQ >= 30** | 27,589,249 | 27,012,845 | 28,295,707 | 43,763,077 | 5,161,279 |
| **# of unique mapped, associated, MAPQ >= 30** | 14,997,809 | 11,799,762 | 21,980,282 | 37,360,138 | 4,622,296 |
| **# of cellular indices** | 1,081 | 841 | 3,184 | 3,743 | 1,291 |
| **Median read count per cellular index** | 9,274 | 8,335 | 4,137 | 6,270 | 2,146 |
| **# of cells, post-filtering** | 975 | 766 | 2,900 | 3,500 | 1,175 |
| **Median read count per filtered cell** | 9,186 | 8,390 | 4,249 | 6,479 | 2,138 |
| **Median *cis:trans* Ratio Across Cells** | 4.43 | 4.34 | 3.69 | 3.42 | 3.96 |

Before deconvolving the resulting data to single cells, we examined the overall distribution of ligation junctions ( *i.e.* contacts). Encouragingly, there were very few contacts between mouse and human (ML1: 0.006%; ML2: 0.008%), demonstrating minimal cross-talk between cellular indices, and that nuclei remain intact through all ligation steps (confirmed through phase-contrast microscopy; **Supplementary Figure 1**). We also examined the *cis:trans* ratio, defined here as the ratio of long-range ( *i.e.*>20 kb) intrachromosomal contacts to interchromosomal contacts (Figure 1c), and found it to be on par with expectation for high-quality Hi-C datasets (ML1: 4.41; ML2: 4.38).

We next split the Hi-C data by cellular index and characterized the number of unique read-pairs associated with each, the vast majority of which should correspond to single cells. When examining a histogram of unique index occurrences as a function of read depth, we noted a bimodal distribution, reminiscent of patterns seen in scATAC-seq datasets^10^, where low-coverage indices likely represent ‘noise’ consequent to tags from free DNA in solution (**Supplementary Figure 2**). After discarding these, we infer 1,081 cellular indices in ML1, with a median of 9,274 unique read-pairs per index (ML2: 841 cellular indices; median of 8,335 unique read-pairs per index). Importantly, we also observe minimal barcode bias across replicate experiments (**Supplementary Figure 3**), as well as similar median *cis*:*trans* ratios per cell (ML1: 4.43 with median absolute deviation (MAD) of 1.66; ML2: 4.34 with MAD of 1.66) (Figure 1d, Supplementary Figure 4).

**Figure 2:**
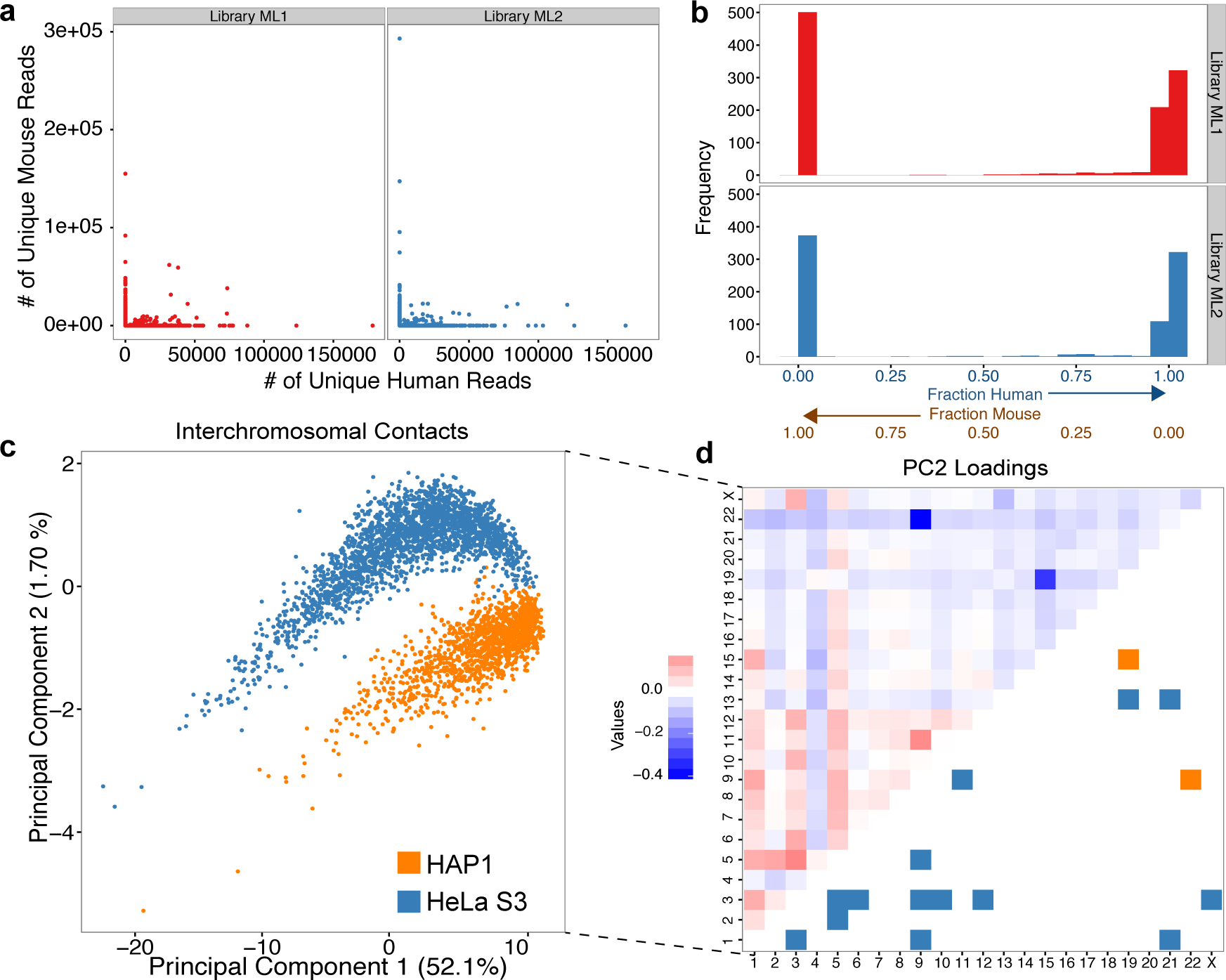
The large number of cellular indices generated through combinatorial single cell Hi-C are overwhelmingly species-specific, and can be separated by cell type. a.) Species mixing experiments enable explicit definition of the “collision rate” in a combinatorial single cell Hi-C experiment. In libraries ML1 and ML2, similar levels of collision (defined as any cellular index with at least 1,000 unique reads, but <95% species purity) are observed, and fall within the expected range. In these plots, points close to the *x* and *y* axes likely represent single cells. b.) Species contamination visualized as a histogram of the fraction of reads mapping to the human genome. Cellular indices (here filtering out all indices with fewer than 1,000 unique reads) are largely species specific. c.) Projection onto the first two principal components from PCA analysis of interchromosomal contact matrices results in separation of HeLa S3 and HAP1, two karytoypically different cell lines ( *n =*3,609 cells). Percentages shown are the percentage of variance explained by each plotted PC. d.) Principal component 2 loadings represent the contribution of each feature (interchromosomal contact) to the observed cell type separation. Strongly blue features appear to be HAP1 specific, while red features appear to be HeLa S3 specific. Known translocations for each cell type are mirrored against the loading heatmap.

The only previously published example of single-cell Hi-C data suggests that high single cell *cis*:*trans* ratios are a hallmark of high-quality single-cell data^9^. The high *cis:trans* ratios that we observe are comparable to those of the 10 single-cell maps generated in that study, which reported a median value of 6.26 (MAD = 0.74), calculated as the ratio of *all* intrachromosomal contacts to interchromosomal contacts ( *i.e.* with no cutoff for minimal intrachromosomal distance). Reanalyzing our own data using this more liberal criterion yielded similar ratios of 6.17 (ML1; MAD = 1.99) and 5.96 (ML2; MAD = 1.94). Of note, our ratios are calculated over 1,922 cellular indices (ML1 and ML2 combined), 857 of which have more than 10,000 unique contacts, compared to the 10 previously reported single cells each with at least 10,000 unique contacts. This comparison illustrates the scalability of combinatorial methods, as compared with methods relying on the physical isolation and serial processing of each single cell.

We designed our experiments to facilitate validation of the single-cell origin of each cellular index. Uniquely tagged cells should be associated with species-specific cellular indices in mixture experiments, with a collision rate broadly defined by a formulation of the “birthday problem^10^.” Consistent with the expected collision rate, we observed that 4.53% of all ML1 cellular indices (4.40% in ML2) were “collisions” (*i.e.* had less than 95% of reads mapping to either the mouse or human genome) (Figure 2a,b). For further analyses we filtered out any cellular indices failing this criterion, while accepting that we remain blind to “within species” collisions. We also filtered out indices where the associated *cis*:*trans* ratio was less than 1 (1.94% of indices in ML1; 1.62% in ML2), which could suggest broken nuclei.

Before continuing, we combined filtered data from ML1 and ML2 with equivalently filtered data from secondary experiments (PL1 and PL2) ( Table 1, Supplementary Figure 5). We then employed a conservative genotype filter^16^ which removed 20.4% of human cellular indices (Supplementary Figure 6), leaving us with a combined dataset of 3,609 human single cell Hi-C maps. Together with mouse data (which were filtered for coverage, *cis*:*trans* ratio, and species purity), a total of 8,141 single cell Hi-C maps were generated across these four experiments.

We next explored whether cell types could be separated *in silico* on the basis of single-cell Hi-C signal. We generated matrices where rows represent single cells, and columns represent the number of contacts between pairs of chromosomes ( Supplementary Figure 7a). Principal components analysis (PCA) on this matrix resulted in separation of single HeLa S3 and HAP1 cells (Figure 2c), which was validated by our programmed barcode associations. Principal component 1 (PC1), which strongly correlated with coverage (Supplementary Figure 8), accounted for the majority of the variance (52.1%), while the combination of PC1 and principal component 2 (PC2; 1.07% of the variance) separated HeLa S3 and HAP1 cells. We then analyzed the “loadings” of our features in PC2, the axis separating HeLa S3 and HAP1 cells, and found that the strongest loadings recapitulated known translocations specific to HAP1^17^ (namely, translocations between chromosomes 15 and 19, and between chromosomes 9 and 22), while other strong loadings corresponded to documented HeLa S3 translocations^16,18^ (Figure 2d). Repeating these analyses by i.) removing specific interactions from the matrices and repeating PCA (Supplementary Figure 9) ii.) using an alternate feature set (interacting 10 Mb intrachromosomal windows; Supplementary Figures 7b, 10), iii.) separating cells by replicate (Supplementary Figure 11), and iv.) sequencing 908 additional human cells (K562 and GM12878; Library ML3 containing 1,175 cells total; Supplementary Figure 12), all recapitulated cell-type separation to varying degrees, demonstrating that PCA can potentially be used to separate cell types on the basis of Hi-C signal (with the caveat that the separations observed here may be driven by karyotype differences between these cell types).

We next examined the heterogeneity present in single cell Hi-C maps in terms of polymer conformation. We plotted contact probability as a function of genomic distance for 769 single cells, each with at least 10,000 unique contacts (Figure 3a), finding that the pattern of scaling observed for single cells was markedly more disperse when compared to a shuffled control where the assignment of cellular indices to reads are randomized, regardless of species analyzed. We then examined the relationship between single-cell power-law scaling coefficients (Figure 3b), calculated between distances of 50 kb and 8 Mb^19,20^, and single-cell *cis*:*trans* ratios, noting a correlation across four out of five experiments (Figure 3c, Supplementary Figure 13) between high *cis*:*trans* ratios and shallow scaling coefficients. Although beyond the scope of our methodological proof-of-concept, these empirical observations of cell-to-cell heterogeneity in contact probability distributions are likely to be highly useful in constraining computational models of mammalian chromosome conformation.

**Figure 3:**
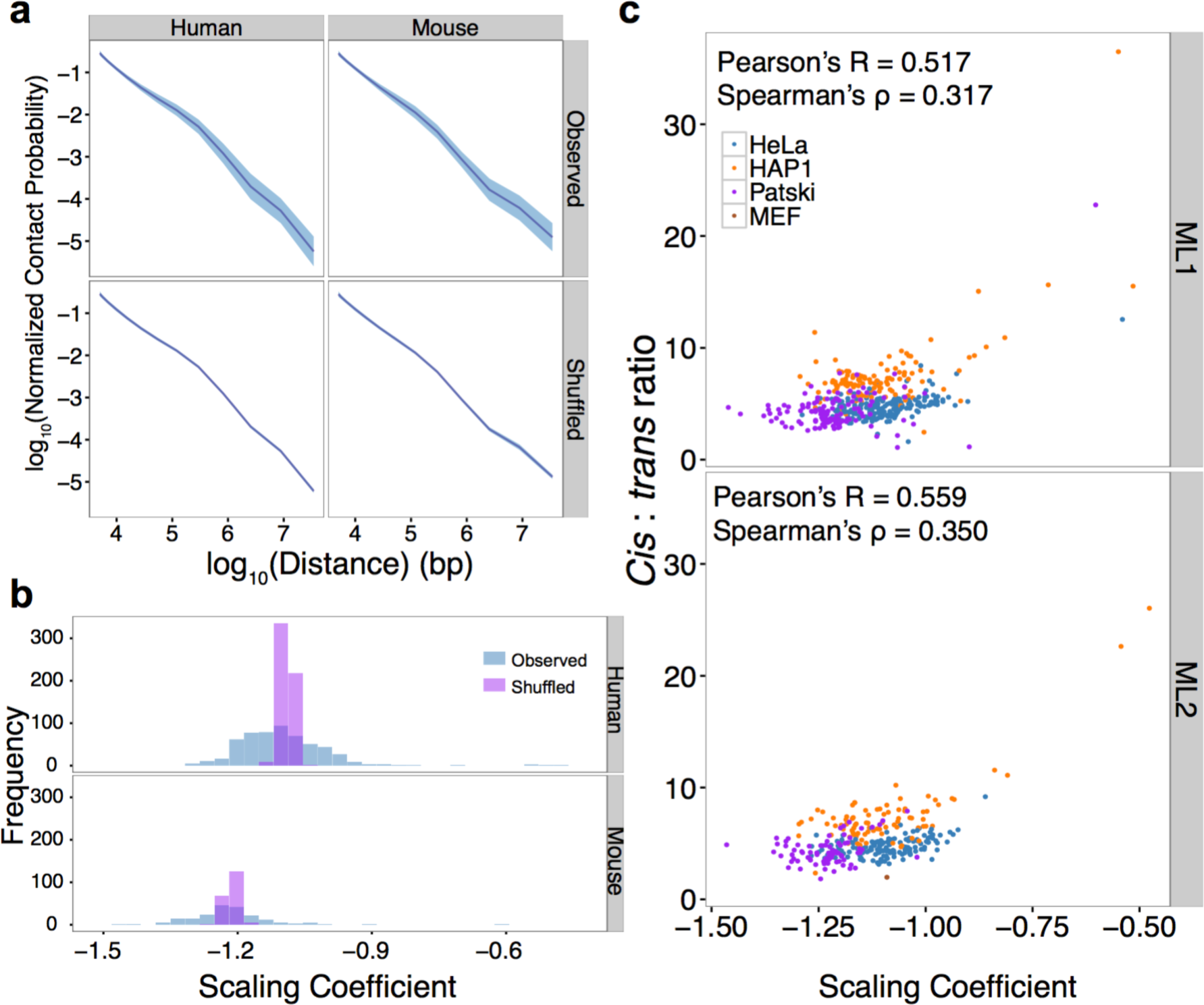
Combinatorial single cell Hi-C captures cell-to-cell heterogeneity masked by bulk measurement, a.) Decay in contact probability for all primary experiment (ML libraries) cells with at least 10,000 unique contacts (*n* = 769 cells). Plotted is the mean contact probability for each bin (purple), along with standard deviation (blue). Shuffled controls where all cellular index assignments have been randomized demonstrate strikingly lower variance compared to observed single cells, for both mouse and human. b) Scaling coefficients calculated for a.), for distances between 50 kb and 8 Mb. Shuffled controls demonstrate a tighter distribution of coefficients compared to the observed single human cells. c.) Single-cell scaling coefficients reproducibly demonstrate positive correlation with single-cell *cis*:*trans* ratios in both mouse and human cells.

In summary, we present a novel method for single-cell Hi-C that relies on the concept of combinatorial cellular indexing for rapid scaling to large numbers of cells. For this proof-of-concept, we applied this method to generate single-cell Hi-C maps for 9,316 cells with at least 1,000 unique contacts. This dataset is two orders of magnitude larger than the only published single-cell Hi-C dataset, with 2,563 filtered cells containing more than 10,000 unique contacts, compared to the 10 existing single-cell maps defined using a similar coverage cutoff. Looking forward, an important technical goal is to further increase the number of unique contacts obtained per single cell, as well as to increase the number of single cells processed per experiment. Importantly, our combinatorial approach is internally controlled in the sense that key steps are carried out in a “single pot”, thus mitigating technical confounders of conventional (serial) replicates of single cell or bulk experiments.

Given the generally similar workflow of our method and traditional bulk Hi-C, it may be possible to incorporate into routine practice, thus adding a ‘single cell’ dimension to Hi-C data production and a means of obtaining single-cell and bulk measurement at once (the latter generated by summing single cells). Furthermore, our demonstration that thousands of single-cell Hi-C maps can be generated in a single workflow, without the need to isolate each cell, demonstrates the power of combinatorial indexing for large-scale single cell biology. Combinatorial indexing may thus be generalizable to additional aspects of single cell or even intracellular biology where DNA barcodes can be incorporated *in situ*.

## Methods

### Cell Culture

HeLa S3 (CCL2.2), primary MEFs, and Patski cells were cultured at 37°C, 5% CO_2_ in DMEM supplemented with 1X Pen-Strep (Gibco), and 10% FBS (Gibco). HAP1 cells were cultured were cultured at 37°C, 5% CO_2_ in IMDM supplemented with 1X Pen-Strep and 10% FBS. K562 cells were cultured at 37°C, 5% CO_2_ in RPMI-1640 supplemented with 1X Pen-Strep and 10% FBS. GM12878 cells were cultured at 37°C, 5% CO_2_ in RPMI-1640 supplemented with 1X Pen-Strep and 15% FBS.

### Cell Fixation

Adherent cells ( *i.e.* HeLa S3, HAP1, Patski, MEF) were washed once with 1X PBS (Life Technologies), trypsinized (0.25% Trypsin-EDTA, Life Technologies), spun down at 500x **g** for 5 min., and resuspended in 20 mL serum-free DMEM (IMDM for HAP1). Cells were crosslinked by adding 1.12 mL (2% final concentration, for HeLa S3, HAP1, and MEF) or 1.4 mL (2.5% final concentration, for Patski) 37% formaldehyde (Alcon) and incubated at RT (25°C) for 10 min., after which crosslinking was quenched using 1 mL 2.5M glycine. Quenched reactions were incubated on ice for 15 min., spun down at 800 **xg** for 5 min., resuspended in 1X PBS, aliquoted into 10E6 cell aliquots, pelleted once again at 800 **xg** for 5 min, decanted, snap frozen in liquid nitrogen, and finally stored indefinitely at −80°C.

Suspension cells (*i.e.* K562, GM1878) were spun down at 500 **xg** for 5 min., resuspended in 20 mL serum-free RPMI-1640, crosslinked with a final concentration of 2% formaldehyde, and processed as above.

### Combinatorial Single Cell Hi-C

For the step-by-step combinatorial single cell Hi-C protocol, see **Supplementary Protocol**. Like the recently published scDNase-seq protocol^21^, combinatorial single cell Hi-C uses carrier plasmid to prevent DNA losses during steps of the protocol where small amounts of DNA are handled.

All oligonucleotide sequences used in this study were obtained from IDT Technologies, and are detailed in **Supplementary File 1**. All libraries were sequenced on a HiSeq 2500.

#### Barcode Programming

Our primary datasets (Library ML1 and biological replicate library ML2), used HeLa S3, HAP1, Patski, and MEFs, with subsets of human and mouse cell types in distinct wells during the first round of barcoding (HeLa S3 + Patski in half of wells; HAP1 + MEFs in half of wells). Our secondary datasets (Library PL1 and biological replicate PL2) were generated using the same cell types, but a subtly different programming scheme (illustrated in Supplementary Figure 14), wherein each well contained only a single cell type during the first round of barcoding. Finally, we generated and lightly sequenced a 5^th^ library (Library ML3), mixing the same murine cell types as before with two new human cell types—GM12878 and K562—in a similar manner to Libraries ML1 and ML2 (GM12878 + Patski in half of wells; K562 + MEFs in half of wells).

### Bridge Adaptor Barcode Design

Bridge adaptor barcodes were drawn from randomly generated 8-mers, such that the following criteria were met: i.) all adaptors must have a minimum pairwise Levenshtein distance of 3; ii.) adaptors must not contain the sequences TTAA or AAGCTT; iii.) adaptors must contain >60% GC content; iv.) adaptors must not contain homopolymers >= length 3; and v.) adaptors must not be palindromic.

### Processing Combinatorial Single Cell Hi-C Data

All code used for combinatorial single-cell Hi-C data analysis will be available (along with all data) upon publication at https://github.com/VRam142/combinatorialHiC. Below, we describe in detail the analytical pipeline used to process combinatorial single-cell Hi-C data. The analytical steps broadly fall under three categories: i.) Barcode Identification & Read Trimming, ii.) Read Alignment, Read Pairing, & Barcode Assocation, and iii.) Cellular Demultiplexing & Quality Analysis.

#### Barcode Association & Read Trimming

First, to obtain round 2 (*i.e.* terminal) barcodes, we use a custom Python script to iterate through both mates, compare the first 8 bases of each read against the 96 known barcode sequences, and then assign barcodes to each mate using a Levenshtein distance cutoff of 2. Reads “split” in this way are output such that the first 11 bases of each read, which derive from the custom barcoded Y adaptors, are removed. Mates where either terminal barcode went unidentified, or where the terminal barcodes did not match, are discarded.

For each resulting “split” pair of reads, the two reads are then scanned using a custom Python script to find the common portion of the bridge adaptor sequence. The 8 bases immediately 5’ of this sequence are isolated and compared against the 96 known bridge adaptor barcodes, again using a Levenshtein distance cutoff of 2. There are cases where the entire bridge adaptor, including both barcodes flanking the ligation junction, is encountered in one mate, and not the other. To account for these cases, we also isolate the 8 bases flanking the 3’ end of the common bridge adaptor sequence (when it is encountered within a read), reverse complement it, and compare the resulting 8-mer against the 96 known bridge adaptor barcodes. Output reads are then clipped to remove the bridge adaptor and all 3’ sequence. Barcodes flanking the ligation junction should match; again, mates where barcodes do not match, or where a barcode is not found are discarded.

The result of this processing module are three files: filtered reads 1 and 2, and an “associations” file—a tab-delimited file where the name of each read passing the above filters and their associated barcode combination are listed.

#### Read Alignment, Read Pairing, & Barcode Assocation

As is standard for Hi-C reads, the resulting processed and filtered reads 1 and 2 were aligned separately using bowtie2/2.2.3 to a Burrows-Wheeler Index of the concatenated mouse (mm10) and human (hg19) genomes. Individual SAM files were then converted to BED format and filtered for alignments with MAPQ >= 30 using a combination of samtools, bedtools, and awk. Using bedtools closest along with a BED file of all DpnII sites in both genomes (generated using HiC-Pro^22^), the closest DpnII site to each read was determined, after which BED files were concatenated, sorted on read ID using UNIX sort, and then processed using a custom Python script to generate a BEDPE format file where 5’ mates always precede 3’ mates, and where a simple Python dictionary is used to associate barcode combinations contained in the “associations” file with each pair of reads. Reads were then sorted by barcode, read 1 chromosome, start, end, read 2 chromosome, start, and end using UNIX sort, and deduplicated using a custom Python script on the following criteria: reads were considered to be PCR duplicates if they were associated with the same cellular index, and if they comprised a ligation between the same two restriction sites as defined using bedtools closest.

#### Cellular Demultiplexing & Quality Analysis

When demultiplexing cells, we run two custom Python scripts. First, we generate a “percentages” file that includes the species purity of each cellular index, the coverage of each index, and the number of times a particular restriction fragment is observed once, twice, thrice, and four times. We also include the *cis*:*trans* ratio described above, and, if applicable, the fraction of homozygous alternate HeLa alleles observed. We use these percentages files to filter BEDPE files (see below) and generate, at any desired resolution, single cell matrices in long format (*i.e.* BIN1-BIN2-COUNT), with only the “upper diagonal” of the matrix included to reduce storage footprint. These matrices are then converted to numpy matrices for visualization and further analysis.

#### Filtration of Cellular Indices

We applied several filters to our resulting cellular indices to arrive at the cells analyzed in this study. We first removed all cellular indices with fewer than 1000 unique reads. We next filtered out all indices where the *cis*:*trans* ratio was lower than 1. Finally, for all experiments we removed cellular indices where less than 95% of reads aligned uniquely to either the mouse (mm10) or human (hg19) genomes. For all human cells from HAP1 and HeLa S3 mixing experiments (Libraries ML1, ML2, PL1, and PL2) further filtration by genotype was performed. For each cellular index, we examined all reads overlapping with known alternate homozygous sites in the HeLa S3 genome and computed the fraction of sites where the alternate allele is observed. We then drew cutoffs to filter out all cells where this fraction fell between 56% and 99%.

We do acknowledge that particular applications (*e.g.* structural modeling) may require more stringent filtration for cellular indices covering single cells. As such, we provide with the raw data a supplementary file specifying the “species purity” of each barcode combination in each sequenced library, along with the number of times DpnII restriction fragments are observed in a cell once, twice, thrice, or four times, with the expectation that given some tolerable noise level, one should only observe restriction fragment copy numbers equal to or less than the copy number of that fragment for that cell type. Relatedly, we note that further inspection of the HAP1 cells used in this study revealed that they were not entirely haploid. HAP1 cells, an engineered haploid line, have faster doubling times compared to HeLa S3, and have been described as having a relatively large frequency of diploid cells^23^. FACS analysis (data not shown) of the stock used for these experiments showed that ∼40% of cells analyzed harbored 2 *n* nucleic acid content, indicating haploid cells in G2 or reverted diploid cells in G1.

### Data Analysis

#### PCA of Combinatorial Single-Cell Hi-C Data

Single-cell matrices at interchromosomal contact resolution (log_10_ of contact counts) and 10 Mb resolution (binarized; 0 if absent, 1 if present) were vectorized and concatenated using custom Python scripts. Concatenation was performed such that redundant entries of each contact matrix (i.e. *C*_*ij*_ and *C*_*ji*_) were only represented once. Resulting matrices, where rows represent single-cells and columns represent observed contacts, were then decomposed using the PCA function in scikit-learn. For interchromosomal matrices, entries for intrachromosomal contacts (*i.e.* the diagonal) were set to 0. For 10 Mb intrachromosomal matrices, all interchromosomal contacts were ignored and all entries *C*_ij_ where |*i – j*| < 3 were set to zero.

#### Calculation of Contact Probabilities in Single Cells

Methods to calculate the scaling probability within single cells were adapted from Imakaev, Fudenberg *et al*^19^ and Sanborn, Rao *et al*^20^. A histogram of contact distances normalized by bin size was generated using logarithmically increasing bins (increasing by powers of 1.12^*n*^). We obtained the scaling coefficient by calculating the line of best fit for the log-log plot of this histogram between distances of 50 kb and 8 Mb. Shuffled controls were generated by randomly reassigning all cellular indices and repeating the above analysis; this importantly maintains the coverage distribution of the new set of simulated “single cells.”

All plots were generated in R using ggplot2 (http://ggplot2.org/).

## Acknowledgements

The authors thank S. Kasinathan, members of the UW Center for Nuclear Organization and Function, and members of the Shendure lab (particularly M. Kircher), for helpful discussions. HeLa S3 cells were used as part of this study. Henrietta Lacks, and the HeLa cell line that was established from her tumor cells in 1951, have made significant contributions to scientific progress and advances in human health. We are grateful to Henrietta Lacks, now deceased, and to her surviving family members for their contributions to biomedical research. Primary MEF aliquots were a gift from Carol Ware. This work was funded by grants from the NIH (5T32HG000035 to VR, DP1HG007811 to JS and U54DK107979 to XD, CMD, WSN, ZD and JS). JS is an Investigator of the 6 Howard Hughes Medical Institute.

## Competing Financial Interests

K.G. and F.S. are employees of Illumina Inc.

**Supplementary Figure 1:**
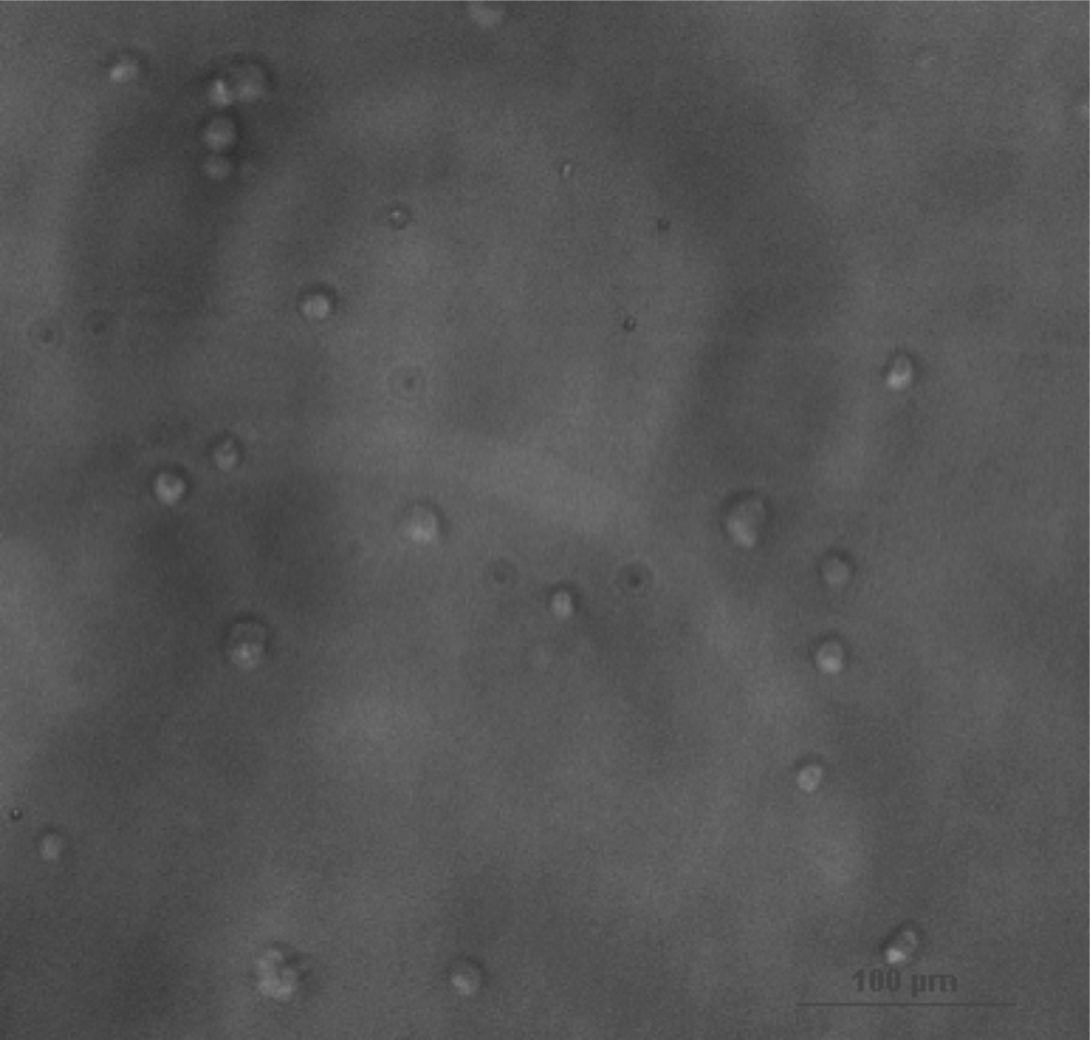
Nuclei remain intact through proximity ligation in the combinatorial single cell Hi-C protocol. Phase contrast microscopy of HeLa S3 and HAP1 nuclei following proximity ligation and serial dilution shows that nuclei remain intact throughout the combinatorial single cell Hi-C protocol (scale bar = 100 um).

**Supplementary Figure 2:**
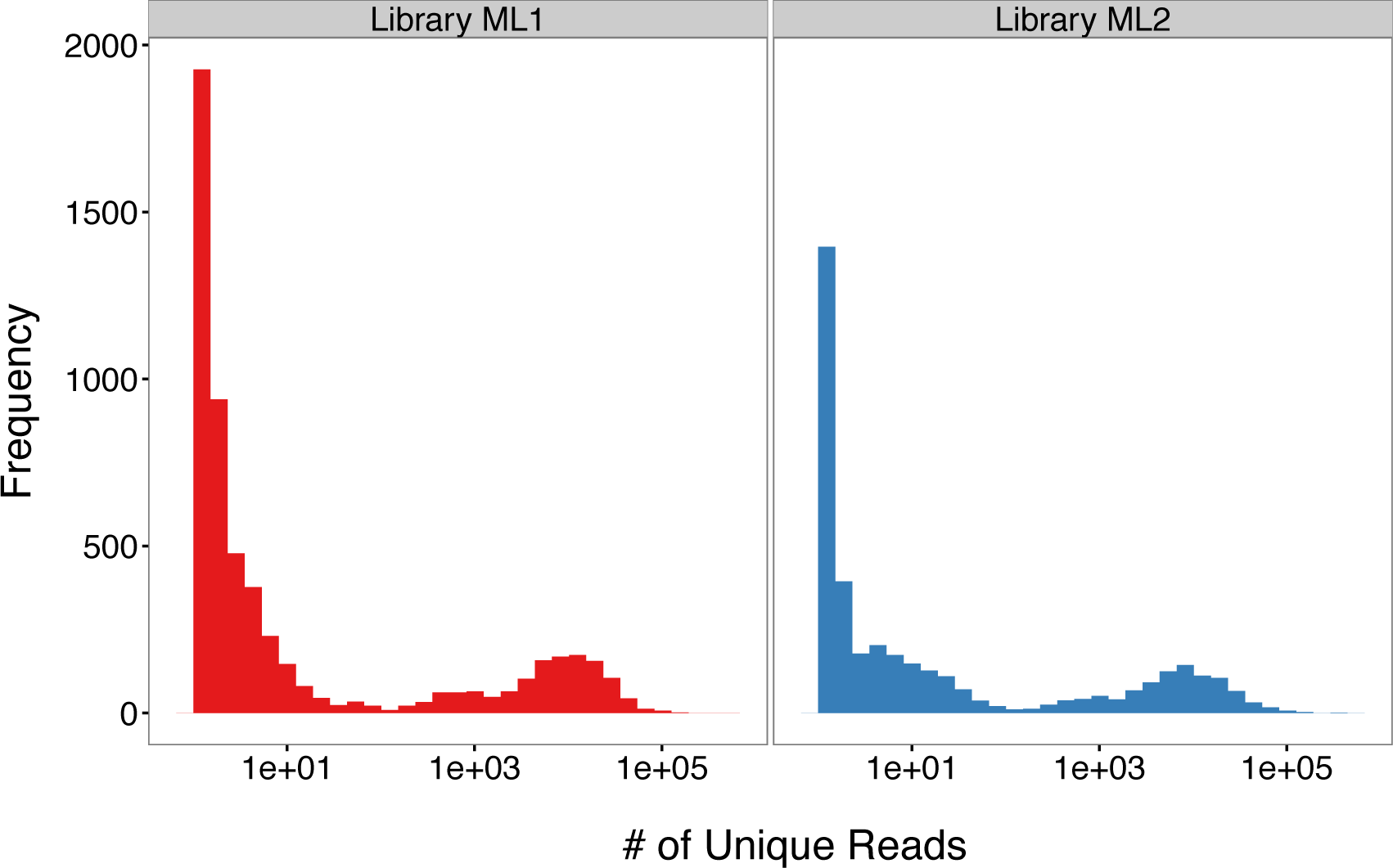
Coverage of combinatorial single cell Hi-C cellular indices follow a bimodal distribution. Examining a histogram of the coverage (*i.e.* ⋕ of unique reads) of combinatorial single cell Hi-C cellular indices in two replicate libraries reveals a bimodal distribution, where low coverage cellular indices likely represent barcoding of free DNA in solution, rather than intact nuclei.

**Supplementary Figure 3:**
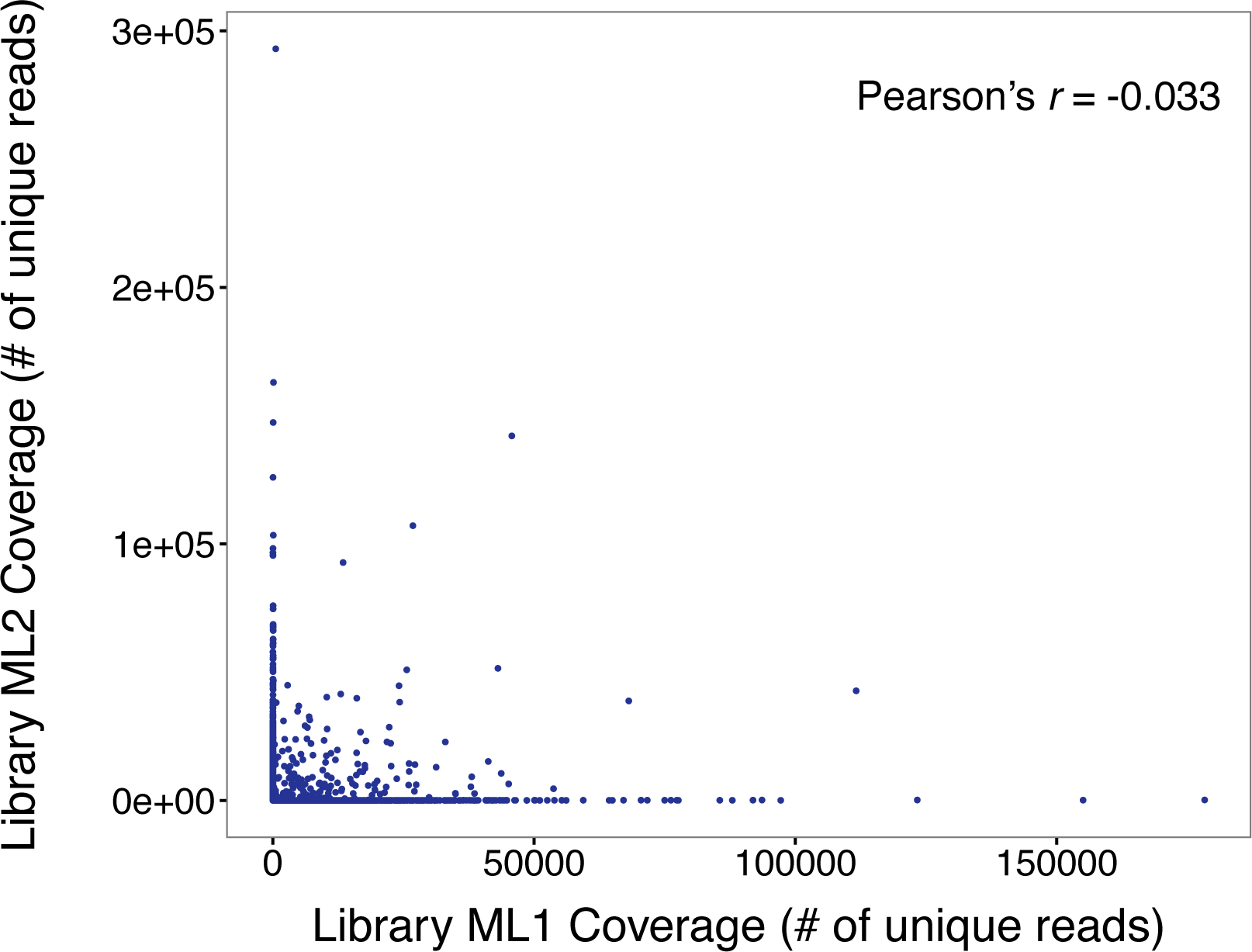
Coverage of cellular indices is not correlated between replicate experiments. Scatter plot of coverage per cellular index for all cellular indices with at least 1 unique read in both replicate combinatorial single cell Hi-C libraries. A Pearson’s r of −0.03 suggests that there is minimal intrinsic bias (*i.e*. “barcode” effect) biasing coverage of particular cellular indices.

**Supplementary Figure 4:**
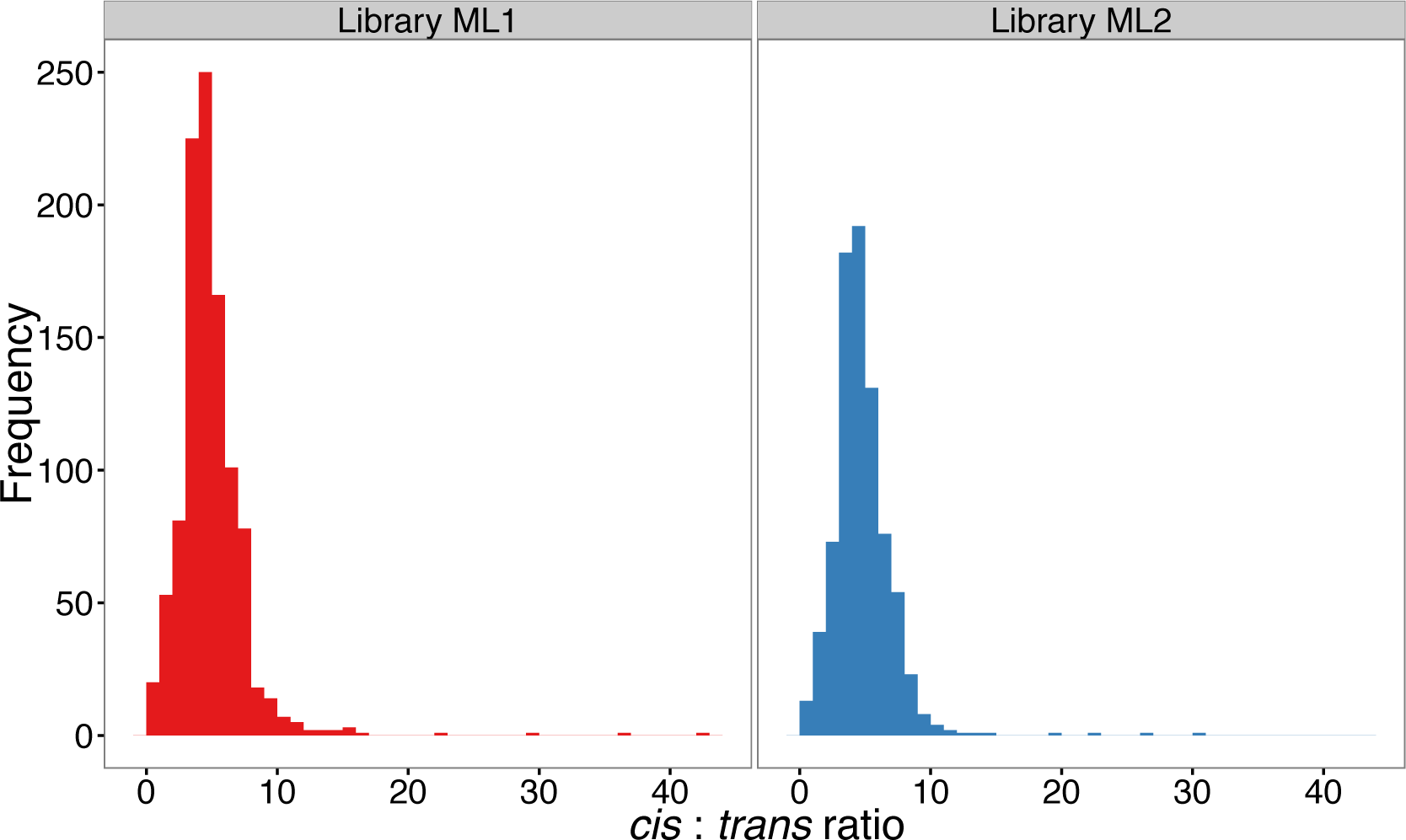
Single cellular indices demonstrate high *cis*:*trans* ratios. Histogram of the *cis*:*trans* ratios for cellular indices over two biological replicates. High *cis*:*trans* ratio suggest that nuclei remain intact during the protocol, and hint at a single-cellular origin for the majority of cellular indices.

**Supplementary Figure 5:**
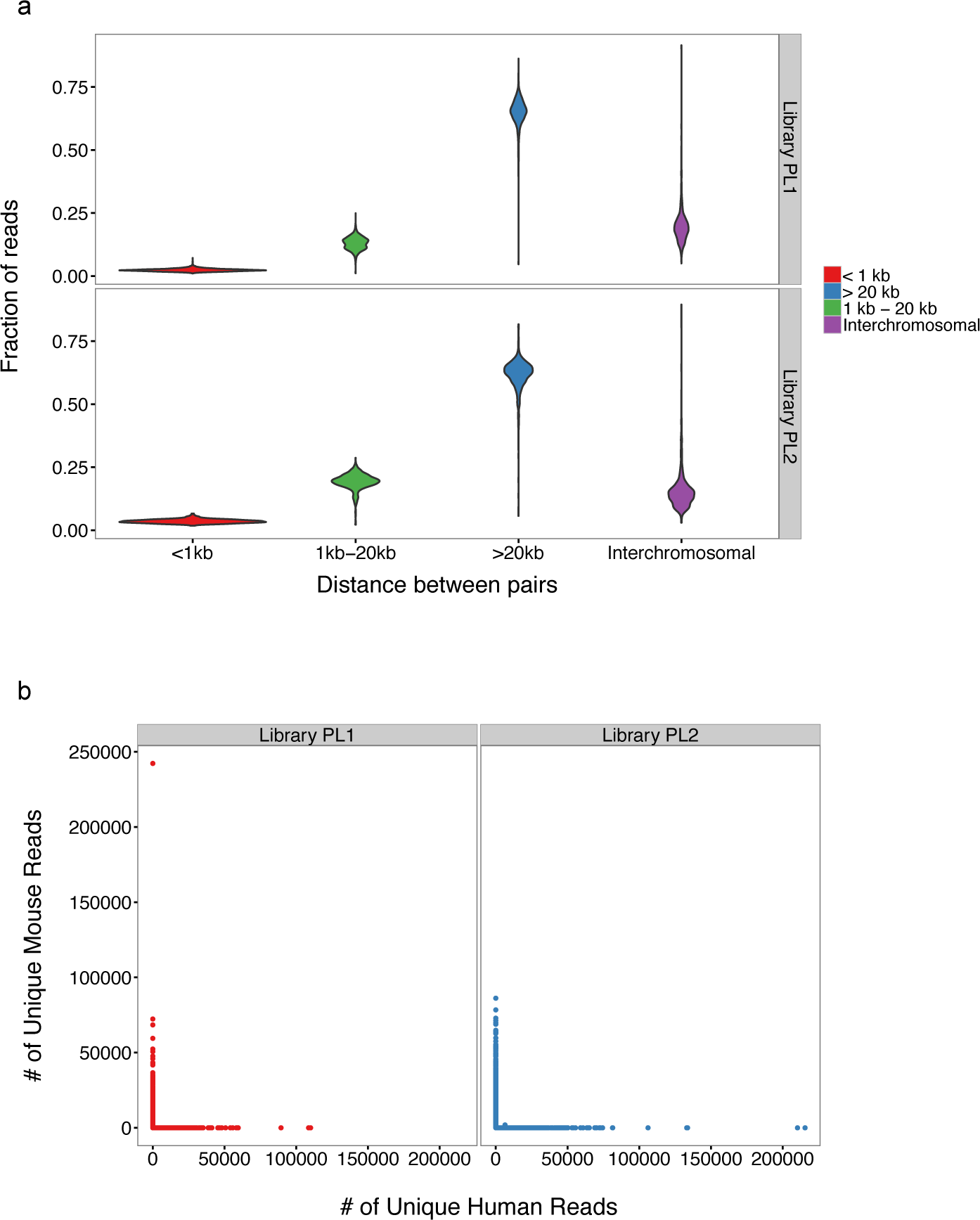
Quality control statistics for PL1 and PL2 libraries are similar to primary experiment libraries. a.) Violin plots showing the distribution of ligation types across all cellular indices with at least 1,000 reads in libraries PL1 and PL2. b.) Species specificity for both libraries.

**Supplementary Figure 6:**
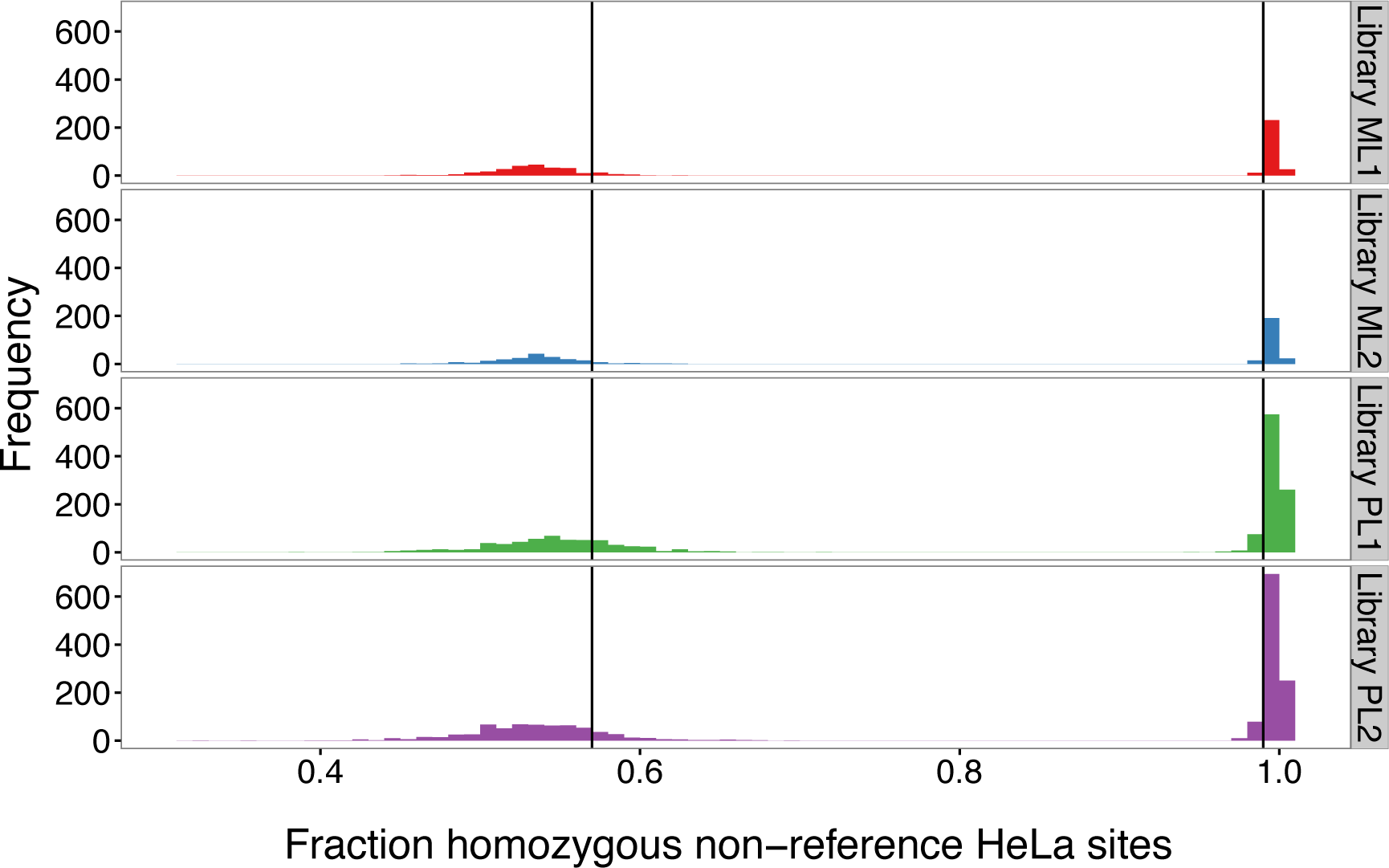
The HeLa genotype enables further filtration of potential barcode collisions in combinatorial single cell Hi-C datasets. We examined all homozygous non-reference sites determined by Adey *et al*^16^ and tabulated the fraction of sites where the non-reference allele was found in our sequencing reads, with the expectation that single HeLa cells should have very high (*i.e.* >=99%) homozygous non-reference alleles at those sites, with reduced fractions indicating contamination by HAP1. For this study, we drew conservative cutoffs of 57% and 99% for each species ( *i.e.* any cellular indices falling between these values were discarded).

**Supplementary Figure 7:**
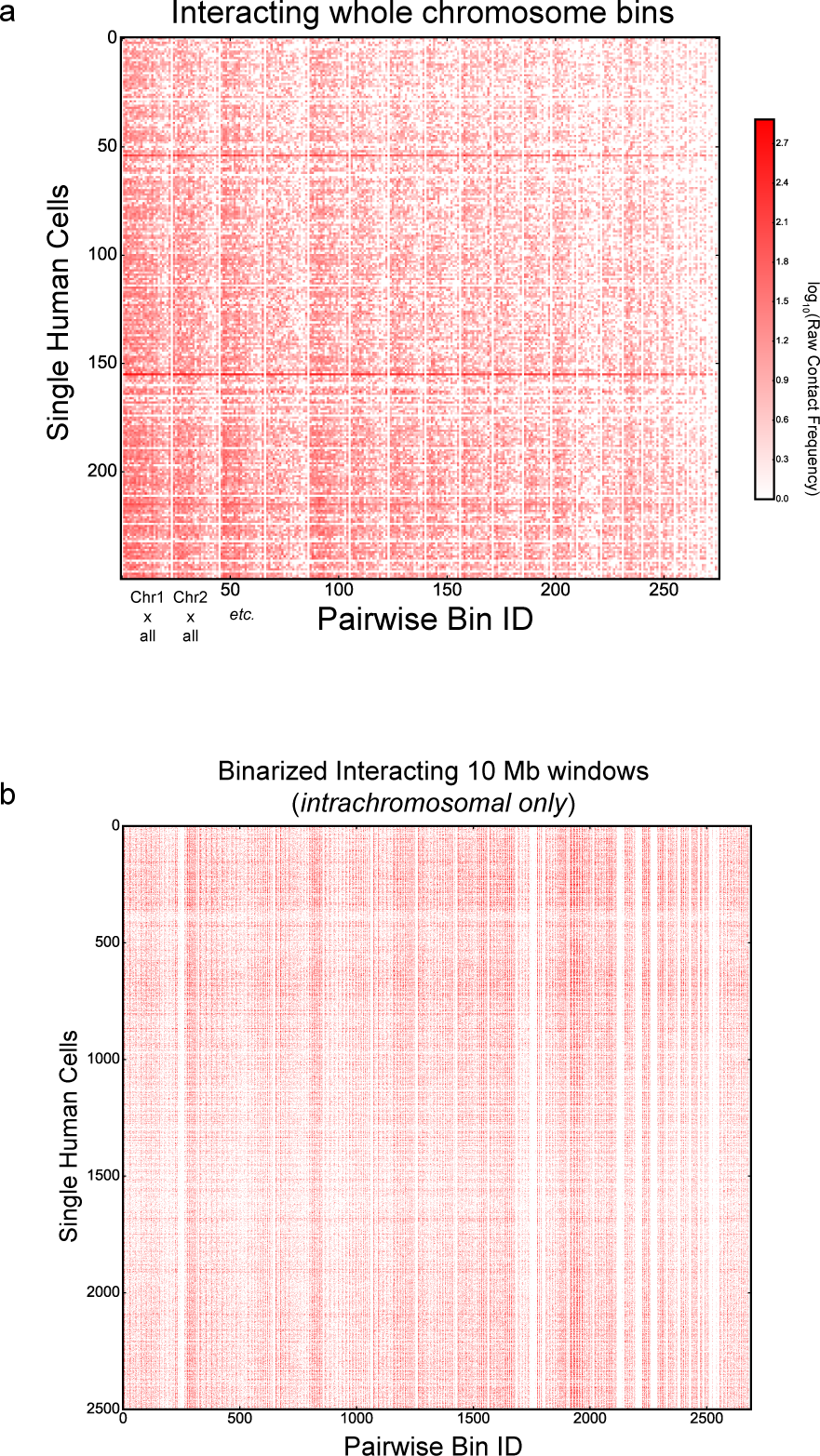
Raw single cell matrices used as input for PCA. a.) A heat map representation of a portion (250 cells) of the input interchromosomal matrix for PCA. Rows represent single human cells, while columns represent pairwise interactions between two whole chromosomes. For this analysis, raw counts were used, and *n* = 3,609 cells. b.) Heat map representation of a portion (2,500 cells) of the input intrachromosomal matrix for PCA. Here, interchromosomal counts were ignored, and interaction frequencies between discrete 10 Mb windows genome-wide were reduced to a binary representation (*i.e.* 1 if present, 0 if absent). Again, *n =*3,609 cells.

**Supplementary Figure 8:**
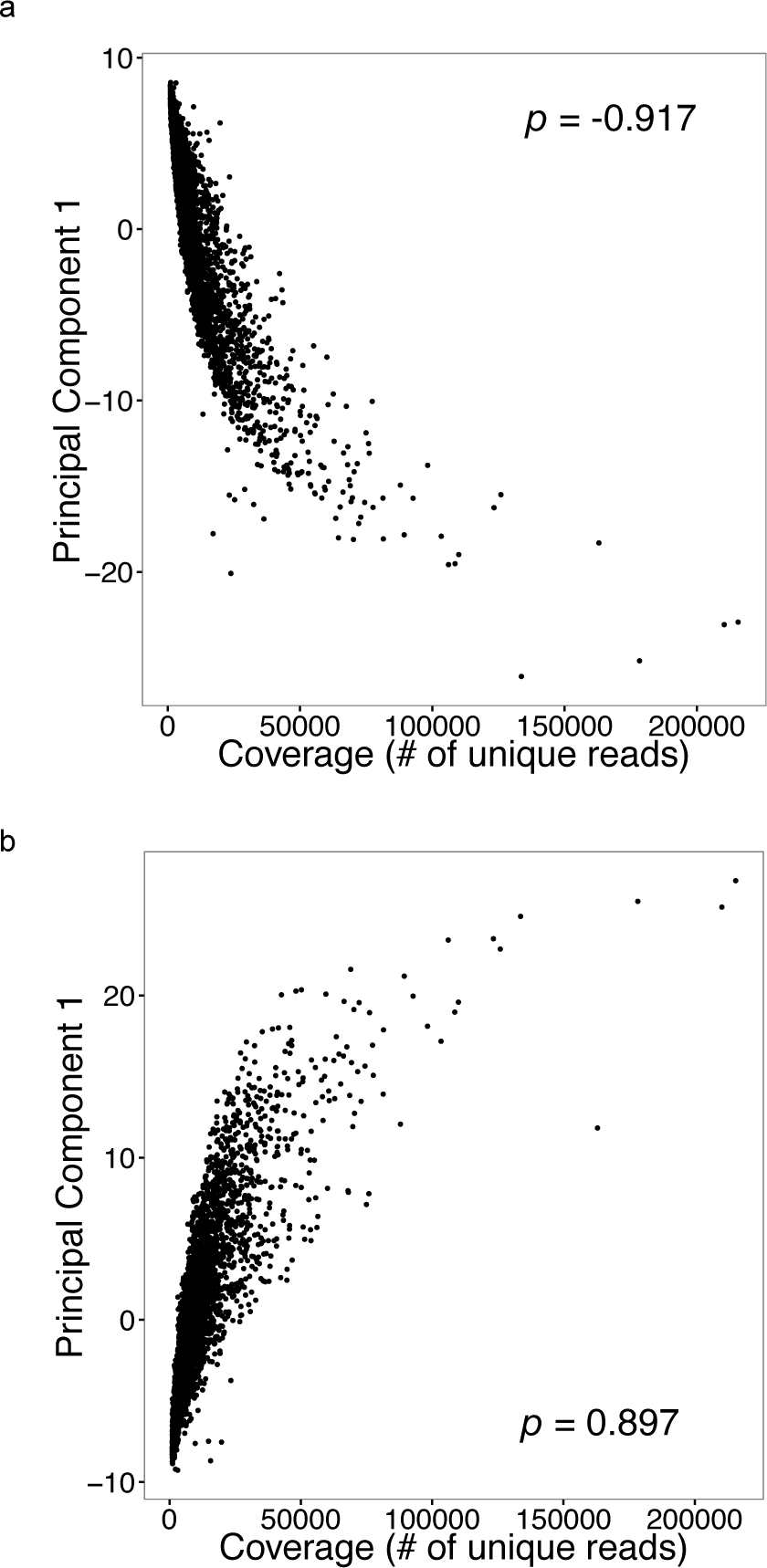
The first component of PCA using both interchromosomal contacts and 10 Mb windowed intrachromosomal contacts strongly correlates with coverage. a.) Correlation between the principal component 1 (PC1) and coverage for interchromosomal interactions (ρ = −0.917). b.) Correlation between the principal component 1 (PC1) and coverage for interacting 10 Mb intrachromosomal windows (ρ = 0.897)

**Supplementary Figure 9:**
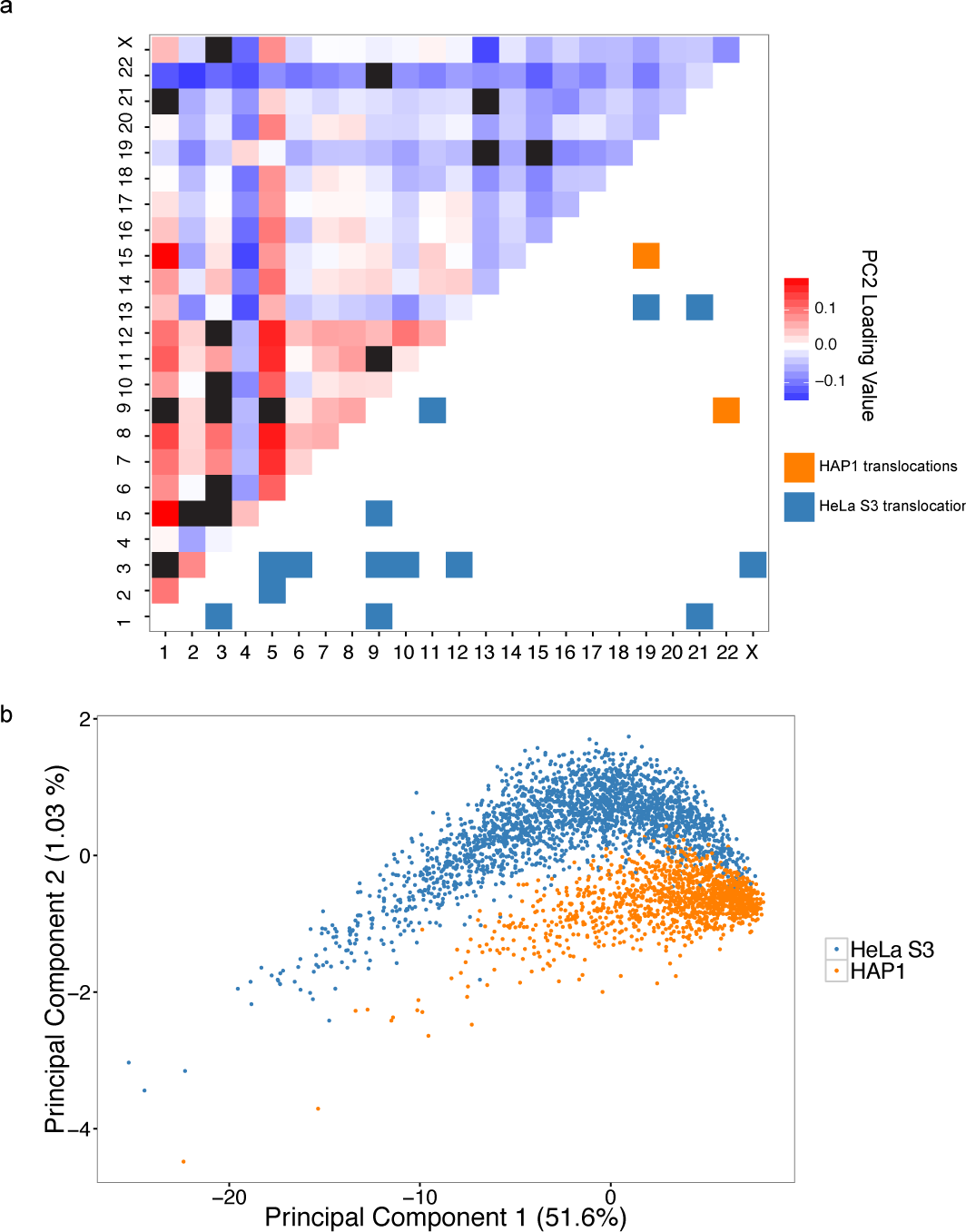
Analysis of principal component loadings for interchromosomal separation experiment reveals that translocations contribute to cell type separation in principal component space. a.) Heat map of loadings for principal component 2 after all known translocations (blacked out entries) are removed from the analysis. b.) After removing all entries corresponding to known translocations from the interchromosomal single-cell Hi-C contact matrix, cell-type separation using PC1 and PC2 is qualitatively worse but still apparent, suggesting that cell-type specific interchromosomal contacts may contribute to the observed separation pattern. Percentages shown are the percentage of variance explained by each plotted PC.

**Supplementary Figure 10:**
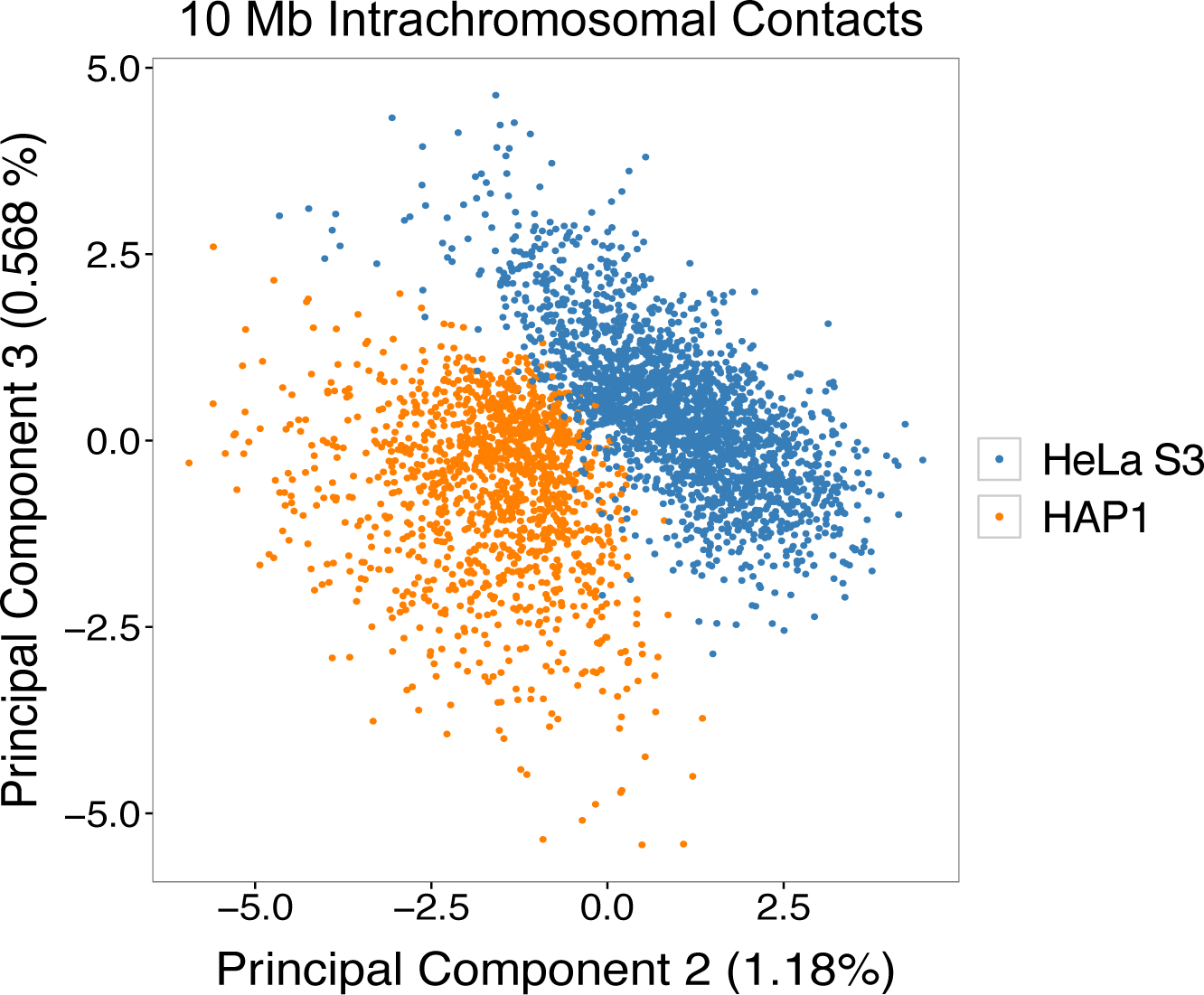
PCA using an alternate feature set still enables separation between HAP1 and K562. Shown is a projection of principal component 2 and principal component 3 from PCA on the intrachromosomal single cell contact matrix (*n* = 3,609 cells). For this analysis, only intrachromosomal contacts between 10 Mb windows were used. The matrix used for this computation is shown in **Supplementary Figure 7b**. Percentages shown are the percentage of variance explained by each plotted PC.

**Supplementary Figure 11:**
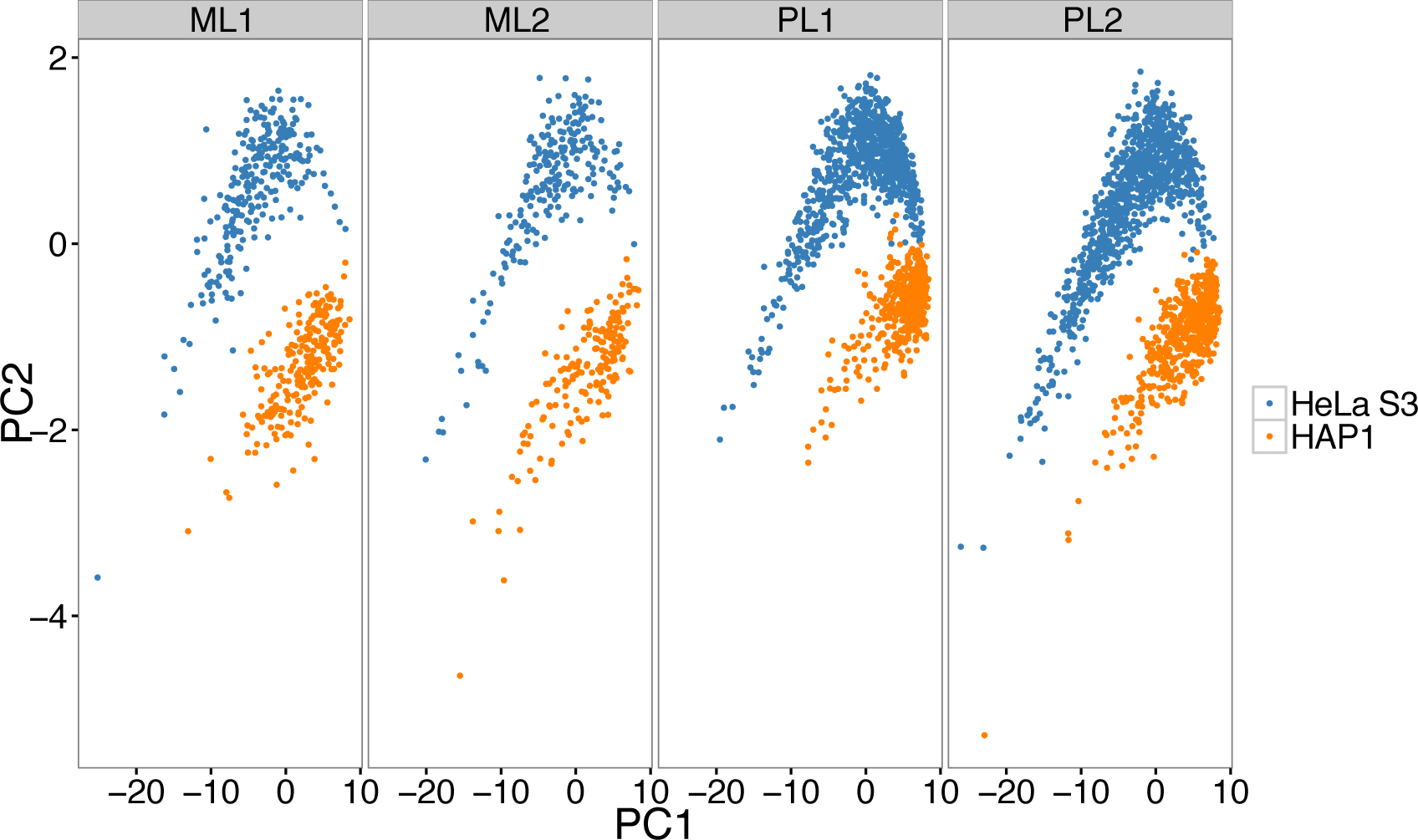
Separation of cell types by PCA is consistent across biological replicate combinatorial single cell Hi-C experiments. Across 4 different libraries, the separation of single HeLa S3 and HAP1 cells is evident, suggesting that this is not simply a technical artifact or batch effect.

**Supplementary Figure 12:**
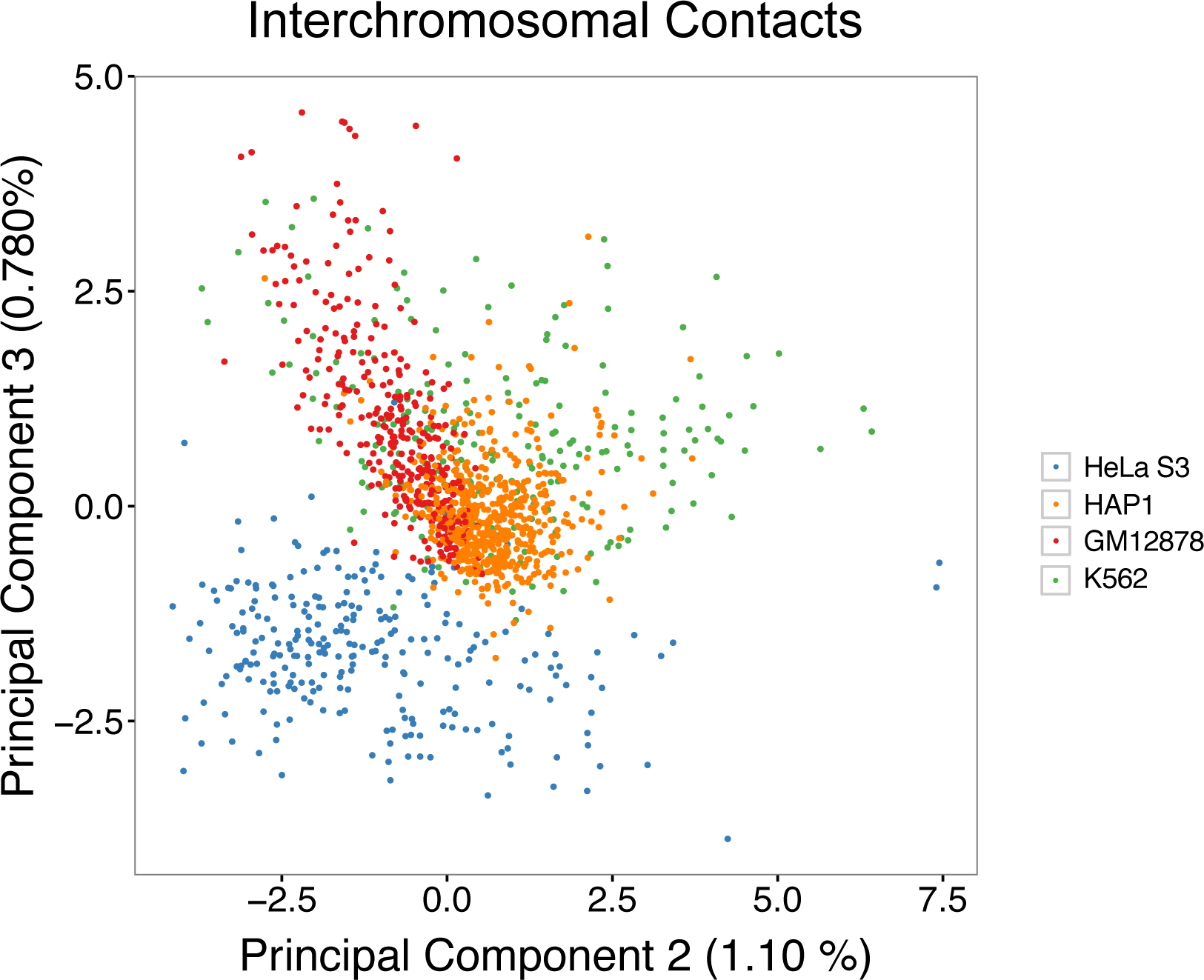
PCA of single-cell interchromosomal contacts using cells from 4 different human cell types results in separation of HeLa S3 from other cell lines. A fifth experiment (Library ML3) containing K562 and GM12878 cells was lightly sequenced and combined with an existing HeLa S3 and HAP1 dataset (Library ML1), resulting in *n* = 1,394 cells. Projection of single cells onto PC2 and PC3 results in separation of HeLa S3 from the remaining three cell types, but weak separation of K562, GM12878, and HAP1. Percentages shown are the percentage of variance explained by each plotted PC.

**Supplementary Figure 13:**
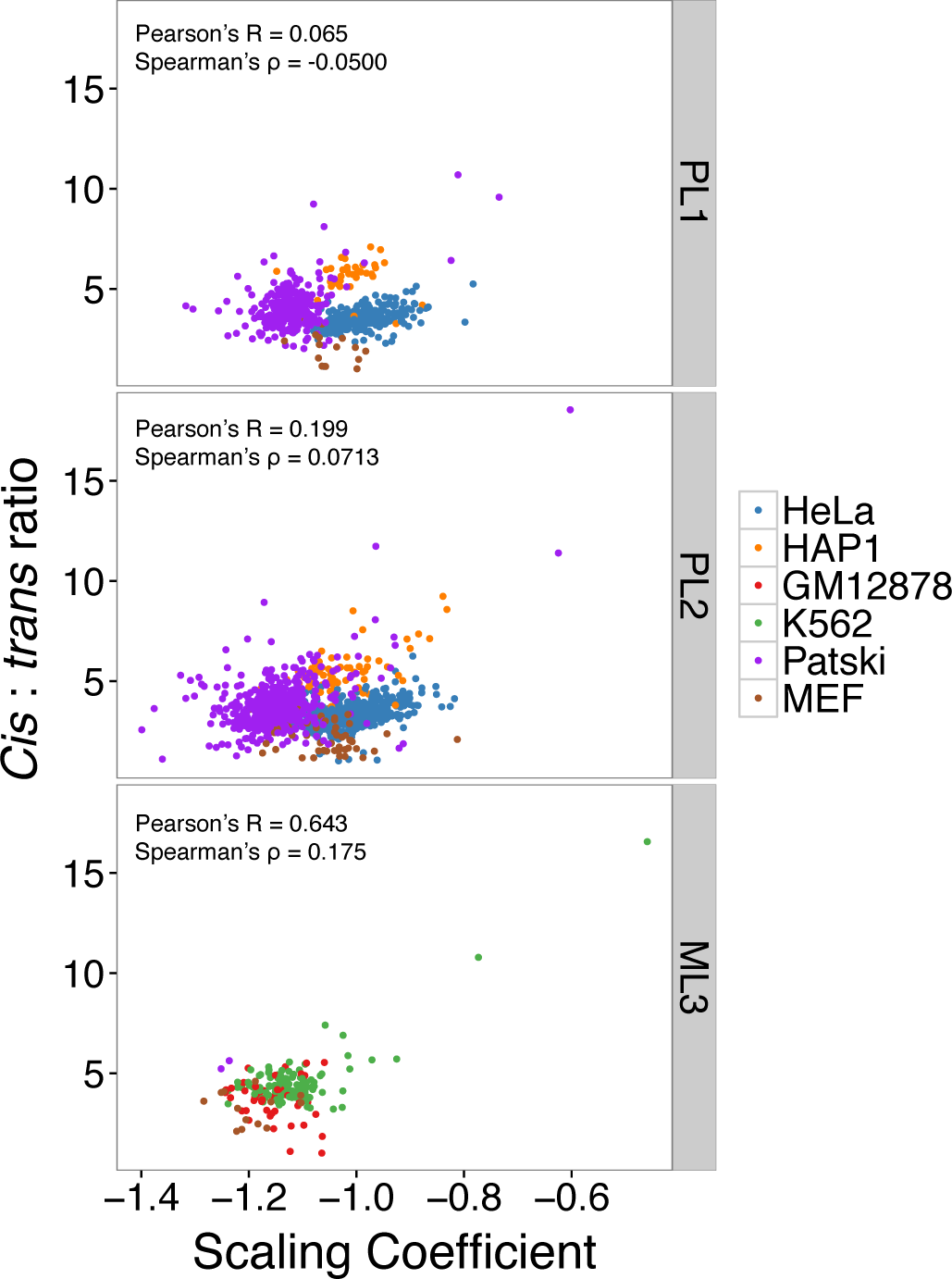
Correlation between single cell *cis*:*trans* ratios and single-cell scaling coefficients is reproducible across combinatorial single-cell Hi-C experiments. We observe a correlation between high *cis*:*trans* ratios and shallow scaling coefficients in both mouse and human cells in both the PL2 (Pearson’s R = 0.199; Spearman’s ρ = 0.0713) and ML3 (Pearson’s R = 0.643; Spearman’s ρ = 0.175) experiments. It is possible that the lack of correlation / weaker correlation shown in PL1 (Pearson’s R = 0.0649; Spearman’s ρ = −0.0500) and PL2, respectively, are a result of shallower sequencing, or sampling (*i.e.* perhaps related to the relative abundance of unsynchronized cells in each phase of the cell cycle).

**Supplementary Figure 14:**
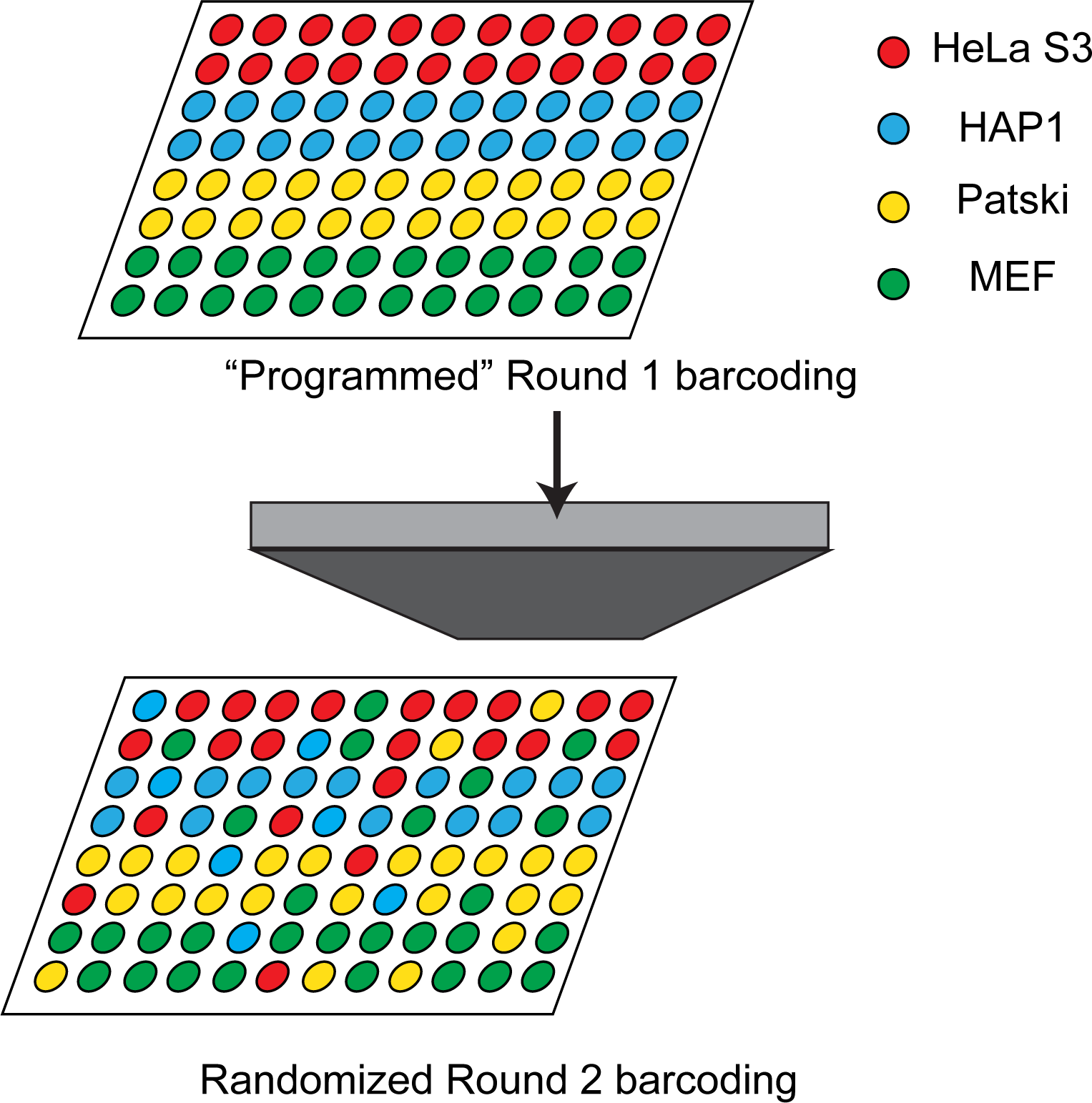
“Programmed” barcoding approaches enable association of cell types with unique first round barcodes. By loading unique cell types into programmed wells during the first round of indexing, we are able to validate cell types *in silico*. This schematic shows how libraries PL1 and PL2 were generated, wherein only one cell type was present per cell. By contrast, for ML1, ML2 and ML3, subsets of wells contained mixtures of one human and one mouse cell type.

